# Improved Functional Classification of Hydrolases through Pairwise Structural Similarity of Reaction Cores

**DOI:** 10.1101/2025.01.09.632218

**Authors:** Sakib Ferdous, Priyanshu Gupta, Yee Chuen Teoh, Karuna Anna Sajeevan, SR Vaishnavey, Kaelie Hainlin, Nigel Reuel, Abir Qamhiyah, Narayana R. Aluru, Manish Kumar, Ratul Chowdhury

## Abstract

We report a systematic pipeline is for extracting the catalytically relevant reactive site in addition to the surrounding allosterically linked residue shells around the reaction site of the most diverse enzyme class - hydrolases, with known experimental structures. We first successfully extract 40196 such hydrolase reaction cores (RC) and collates them into a publicly accessible reaction core collection (RC-Hydrolase). We perform 128M pairwise shape comparison across RC-Hydrolase using a three-dimensional search engine and present 155,329 pair instances clustering them by 60% or higher similarities in a publicly available, visually interactive dataset. Robustness of defined RCs is shown to successfully capture experimentally known function-enhancing mutations distal to the active site in PETases. Allowing comparisons of enzyme reaction centers across functional spaces (ligands bound, EC classification numbers, and expression hosts) enables identification of enzyme backbones which can be minimally mutated to accommodate more than one type of catalytic activity thereby aiding rational design of multifunctional enzymes. We also demonstrate how such versatile enzyme backbones could be leveraged by the latest diffusion-based protein design models to design bespoke libraries of small molecule inhibitors, and structurally stable multifunctional enzyme pockets. With only sporadic successes in multifunctional enzyme design thus far, we provide strong structural priors for machine-learning-guided advanced enzyme engineering in the future.

## Introduction

Multifunctional enzymes play an essential role in various industrial applications and have already been explored as a viable route for enhancing efficiency and sustainability across sectors such as food, pharmaceuticals, and biofuels^1–3^. Enzymes catalyze substrates using a set of active site residues supported by additional degrees of covalently/ non-covalently interacting, neighboring residues (including distal allosterically linked residues) comprising reaction cores (RC hereafter). Reaction cores are pockets or grooves on the enzyme surface centered around key catalytic and substrate/ reaction intermediate stabilizing amino acids that offer the appropriate chemical microenvironment for the substrate(s) and cofactors to bind and product(s) to escape^4^. The rest of the reaction core structure scaffolds around the active site, maintaining the necessary shape and stability. The first step for enzyme engineering applications is to interpret, represent and analyze enzyme metadata. Enzyme metadata exists in different forms - amino acid sequences, whole three-dimensional structures from Protein Data Bank (PDB) entries, and functional kinetic rate parameters^5^. Traditional methods for analyzing and interpreting these data involve sequence alignment, homology sequence/ structure modeling, clustering and molecular docking (of substrates, cofactors, and reaction intermediates) studies^6^. Recent advents in sequencing techniques and machine-learning (ML) based prediction of structure from single-sequence^7^ and multiple sequence alignments ^8^ has augmented the list of high-quality enzyme structures significantly (AlphaFold DB^9^, ESM Atlas^10^). However, the absence of established biochemical ground rules for enzyme design still limits the ability for zero-shot engineering of enzymes with a target functionality ^11^. We provide a new viewpoint to analyze enzyme structure data and provide a comprehensive, visually interactive database to bolster identification of easy-to-engineer backbones in hydrolase enzymes ^12^.

Evolution has fine-tuned natural enzymes to be highly efficient and specific in their role of life-sustaining metabolic and signaling processes ^13,14^. However, when it comes to industrial, ex-vivo applications, baseline activity of natural enzymes does not always meet target, commercial requirements, such as - higher stability, alternate solvent stability, activity under a wide window of operating conditions (temperature and pH-wise), and resistance to proteolytic degradation ^15^ or small-molecule inhibition ^16^. Overall, the focus of enzyme engineering is to identify a stable biocatalyst that can cleave an intended bond in a target molecule (i.e., be of a desired enzyme classification (EC) number) and retain that functionality in an over a wide range of operational conditions ^17^. One possible approach thus far, albeit with sporadic success, has been - grafting catalytic amino acids for a desired reaction onto a stable enzymatic backbone which meets the industrial process requirements ^18^. However, this involves three major challenges – (a) finding suitable candidate protein structures with reactive grooves big enough where the catalytic residues can be grafted to ensure the right side-chain orientation for the desired biochemical reaction, (b) accommodating the rest of the neighboring amino acids to ensure minimal departure from the local geometry of the reactive core, and (c) ensuring that the final enzyme can be expressed in a host organism of interest for facilitating biomanufacturing scale-up. The current work alludes to all these three challenges by performing functional clustering of all known hydrolase enzymes (with reported experimental structures) on the bases of the shape of their reaction cores (with catalytic site and neighboring allosteric shells; **Figure 1**) and provides a computational recipe to design highly/ multifunctional enzymes.

**Figure 1:**
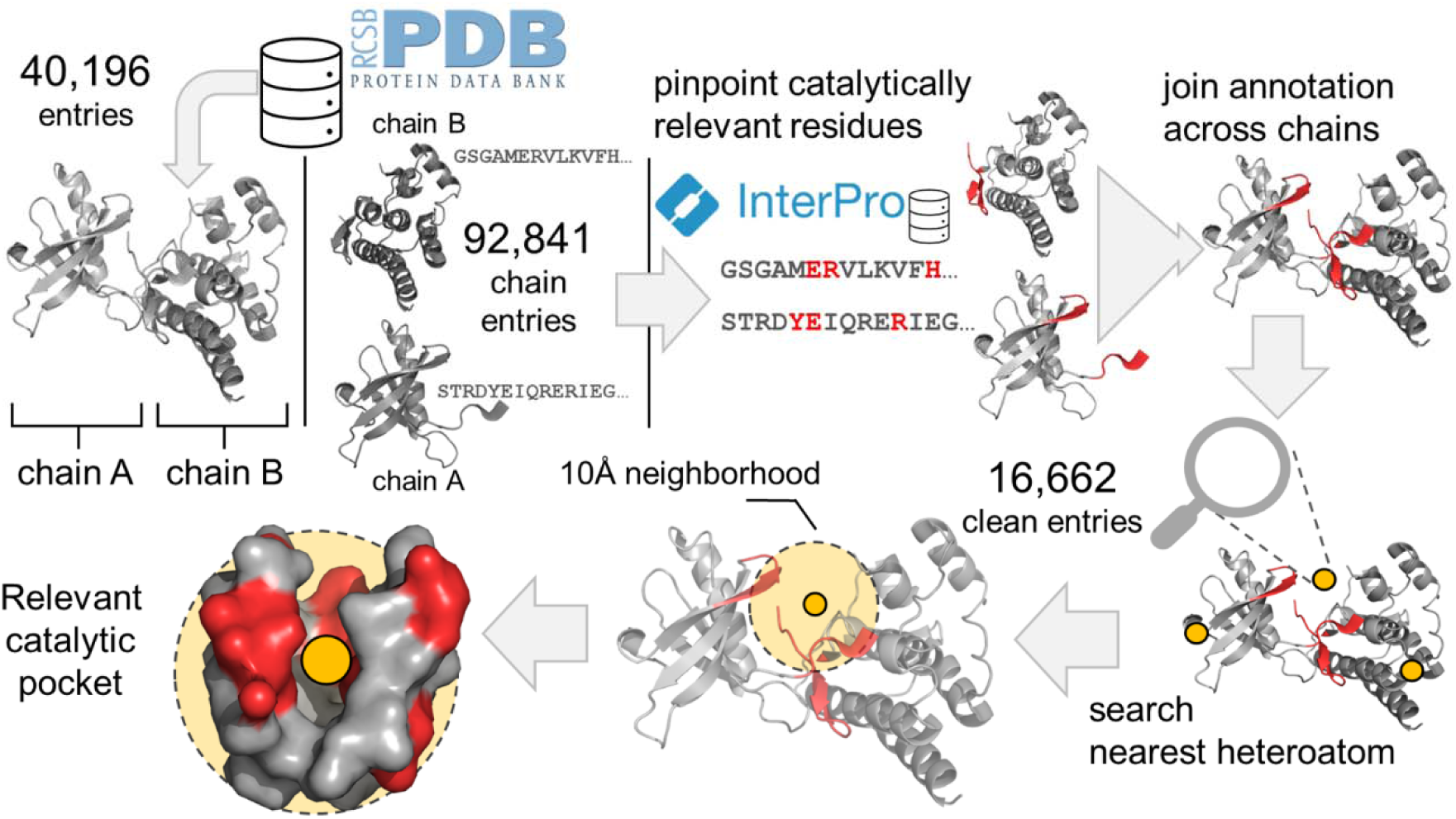
Workflow to get the reaction centers using precise experimental annotations of catalytic residues. 40196 PDB structures are obtained from RCSB databank, the sequences corresponding to each of the chains are extracted, these sequences are sent to Interpro database to identify the conserved domain and functions. The conserved residues are traced back to the structure, with the active site (red) as centroid containing covalently and non-covalently bound resides within 10 Å distance. This functional subsurface of the enzyme defines our reaction core (RC).

Towards the second challenge, rational design efforts were limited by the number of maximum mutations explored as searching the sequence space becomes combinatorically explosive (NP-complete^19^) and experimentally prohibitive. While some successes have been obtained using integer (IPRO^20^+/-) and Monte Carlo sampling (Rosetta3^21^), the recent advent of generative artificial intelligence (AI) to design proteins (RFDiffusion^22^, Genie2^23^) have demonstrated significant successes. Using generative AI, it is theoretically possible to hallucinate the ‘right’ neighboring residues around the newly mutated amino acids wherein the final structure is validated using AlphaFold2. Notwithstanding, the first challenge of geometrically optimal grafting of *de novo* catalytic residues remains elusive to fundamental biochemical/ data-driven rules. Using protein structural metadata and evolutionary deep learning based embeddings (ESM3/ ESMC ^10^) to find a putative enzyme fragment that contains all structural priors to catalysis has not been rigorously elucidated yet. A well-annotated, functionally categorized, query-able database of such key fragment of all known hydrolase enzyme is critical. We have seen accounting for overall sequence and structure information for designing an enzyme is largely lossy as all such encodings are trained to predict protein stability (at best) and not a specific functionality. To this end, in this work, we construct such a sequence-structure database of catalytically relevant RCs for all known hydrolases and a means to cluster them by the shape similarity of only their RCs. RC-Hydrolase is hosted on rc-hydrolase.onrender.com.

### Clustering enzymes by sequence homology

There is a significantly higher number of enzyme sequences than experimentally determined structures. Consequently, the most popular and feasible way to assess and analyze enzymatic proteins is to homology-based alignment (for example - pBLAST^24^) and clustering (for example MMseq2^25^ and Clustal Omega^26^). The key assumption is that-similar sequences fold into similar structures and thus perform similar functions. In practice, however, that is not always the case and there are instances where similar sequences and structures exhibit different functions or different sequences/structures exhibiting same functions because they operate in different cellular contexts ^27 28^. For example – myoglobin and hemoglobin share about 93% sequence similarity but they have differences in their quaternary structure ^29^. Actin, a protein critical for cell structure and motility, although does not have a high degree of sequence similarity, shares a surprising degree of structural similarity with the Hsp70 family of heat shock proteins, which function as molecular chaperones ^30^. Building on biocatalytic intuition, if three amino acids belonging to a catalytic triad of an enzyme are mutated (to alanines), the protein will not be able to perform any function, desptite retaining high sequence and structural similarity. The location of an amino acid within a protein’s three-dimensional structure can thus significantly affect its role and importance. A residue outside the active site is seemingly unimportant for enzyme activity but often are found to be crucial for allosteric regulation, affecting the protein’s activity in response to binding and release kinetics of substrates, reaction intermediates and products from far away ^31^.

### Clustering enzymes by overall structure

With the emergence of protein-based large language models (ESM2 ^32^, AminoBERT ^33^), transformer networks (Evoformer of AlphaFold2 ^34^, ESMFold ^35^, AI-based prediction of enzyme structures has opened doors towards incorporating 3D-structural information along with sequence to cluster proteins with higher functional fidelity. Available information used to train these models include experimentally confirmed structure and sequence combinations and inter-residue co-evolutionary dependence. So, sequence-based structure or function prediction is not free from bias ^36^. Given the new paradigm of ML-based structure predictions, researchers have developed clustering algorithms which can group not-yet-crystallized/ predicted 3D structures of entire proteins towards understanding functions, for example – FoldSeek ^37^.

Regardless, clustering enzymes by their entire structures have their own caveats. Essentially, as previously mentioned - the function of the enzyme depends on a very focused chemical sub-surface (the reaction core) in the enzymatic fold. The rest of the enzyme’s bulk evolves to hold the catalytic center and the allosterically linked neighboring shells (RC) in the right orientation, and even absorb evolutionary and/ or solvent-stabilizing mutations (surface residues) without undergoing structural change to the catalytic reaction core. In other words, the action of the enzyme depends on how the primary (two/ three) cataytic residues are oriented. The diversity in the rest-of-the protein structures pose a significant challenge, as much of the structure is not directly informative about the biochemical reactions being catalyzed. Consequently, clustering these diverse structures suggests numerous post-processing and fine-tuning, complicating the selection of ideal candidates for engineering ^38^. Also, for special cases of natural enzymes such as extremophiles, the sequence is largely unrelated evolutionarily to the known animal kingdom, rather originating in a specific extreme ecological niche. As these enzymes cannot be characterized by homologous analysis, an alternate, catalytically focused scheme is likely to help analyze the functions of such enzymes which will consider the reaction core’s folding for the catalytic action. We hypothesize that the real structural prior between proteins of related function is neither the sequence nor structure, but the reaction core. Thus, it makes more intuitive sense to cluster the reaction core structures only for a high-fidelity functional clustering. A publicly available database of reactive cores is expected to enable focused functional comparison across multiple enzymes of similar/ different biochemical functions conferring a structural basis to designing highly/ multi-functional enzymes. We validate this claim by demonstrating how two enzymes with different substrate preferences, but high RC similarity (RCSim) can be used as structural anchors to design novel multi/ high-functional enzymes.

### Clustering reaction cores (RCs)

Traditional algorithms for comparing protein grooves (i.e., catalytic pockets) include geometry-driven methods like clique detection, triplet and quadruplet matching, pairwise and template-based alignment and more recently machine learning methods. It is noteworthy that most of the existing databases for this purpose are currently out of maintenance and need additional coding to execute them and most importantly none of them come with a publicly accessible website or locally installable codes/ GitHub repositories ^39^. A visually, interactive, publicly available database for matching reaction cores with high structural similarity across different enzyme classes (functions) for a user preferred expression host organism is also absent ^39^. Although it is possible to mine such information from the PDB, they do not offer a way to compare catalytically relevant parts of different enzyme. Structure comparison in PDB is limited to whole-structure comparison only. Some of the reasons for the paucity of focused structural databases can be attributed to the cost of maintaining the database and the challenge of updating them frequently. In contrast, the sequence-based databases are rich, robust, regularly maintained and updated. Regardless, we have seen how 3D structure databases of proteins (AlphaFold DB^9^, PDB^40^, ESM Atlas^10^) have enabled multiple, powerful, machine learning efforts in the last few years ^7,8^. Protein subsurface is also discussed in literature for the purpose of comparison, classification and identification ^41^ without any emergent, downstream design strategy or publicly accessible visual repositories so far. Complementing the existing structure databases with such a reaction core shape matching platform can leverage novel generative AI and structure data towards the goal of designing novel enzymes. To this end, a beta-version of RC-Hydrolase has been already approved to be listed in the PDB from the *Structure Classification and Analysis* page of *Additional Resources* (available from: https://www.rcsb.org/docs/additional-resources/structure-classification-and-analysis).

In this work we outlined a computational protocol for extraction of reaction cores (**Figure 1**) starting with all known hydrolase enzyme structures using precise experimental annotations of catalytic residues aggregated in Interpro^42^. We extracted reaction cores of 16,036 hydrolase structures spanning six major EC functional sub-class of interest – 3.1 through 3.6 (namely – esterase, glycosylase, etherase, peptidases, non-peptide C-N bonds breakers, and acid anhydrides) across four domains of life (i.e., expression hosts) namely archaea, eukaryotes, and bacteria. We also clustered the reaction core structures using a custom shape mapping algorithm (discussed in Methods). Using the structural clusters, we constructed a visual, interactive, publicly available database (RC-Hydrolase) for matching reaction cores with high structural similarity across different enzyme classes (functions) and host organisms. The RC-Hydrolase dataset (available through: https://rc-hydrolase.onrender.com/) contains all pairwise RCSim scores to describe the structural similarity of RCs in all hydrolases. This can also be extended to include predicted enzyme structures. We further described a possible computation protocol to leverage RC-Hydrolase with generative AI based protein design tools, such as ProteinMPNN, to design multifunctional hydrolases.

## Results and Discussion

### Reliability of reaction core definition to capture effects of distal, allosterically linked mutations

Prior studies^43^ focused on using only the catalytic groove of an enzyme to identify beneficial mutations have shown that majority of these mutations are clustered outside the catalytic site but within 15Å of active site^43^. This indicates enzymes are largely intolerant to multiple mutations directly at the catalytic site - but select distal mutations can regulate catalytic activity through a short/ long-range electrostatic and hydrophobic packing network (allostery). We intend to show through two industrially relevant enzymes that our definition of reaction core (with additional allosteric shells around the catalytic groove) is sufficient to capture known beneficial mutations in these enzymes.

First, in a recently reported PET-consuming microbe *Ideonella sakaiensis*, an engineered PET hydrolase mutant (PDB accession id: 5XH3) when expressed in *E. coli* demonstrated 5.5-to 120-fold higher activity than wild type. Using the said PET hydrolase ^44^, we illustrate that our reaction core (RC) definition includes all the relevant catalytic residues, as well as engineered ones (**Figure 2**). This alludes to the generalizability of reactive core definition to relevant hydrolases enzymes (outside the RC-Hydrolase database) to be able to include catalytically critical residues distal from the active site. Recent experimental data on enhanced PET hydrolase catalysis upon mutation of Asp83 and Asp89 demonstrates that RC sub-surfaces are sufficient to capture all relevant mutational loci for engineering enhanced turnover.

**Figure 2.**
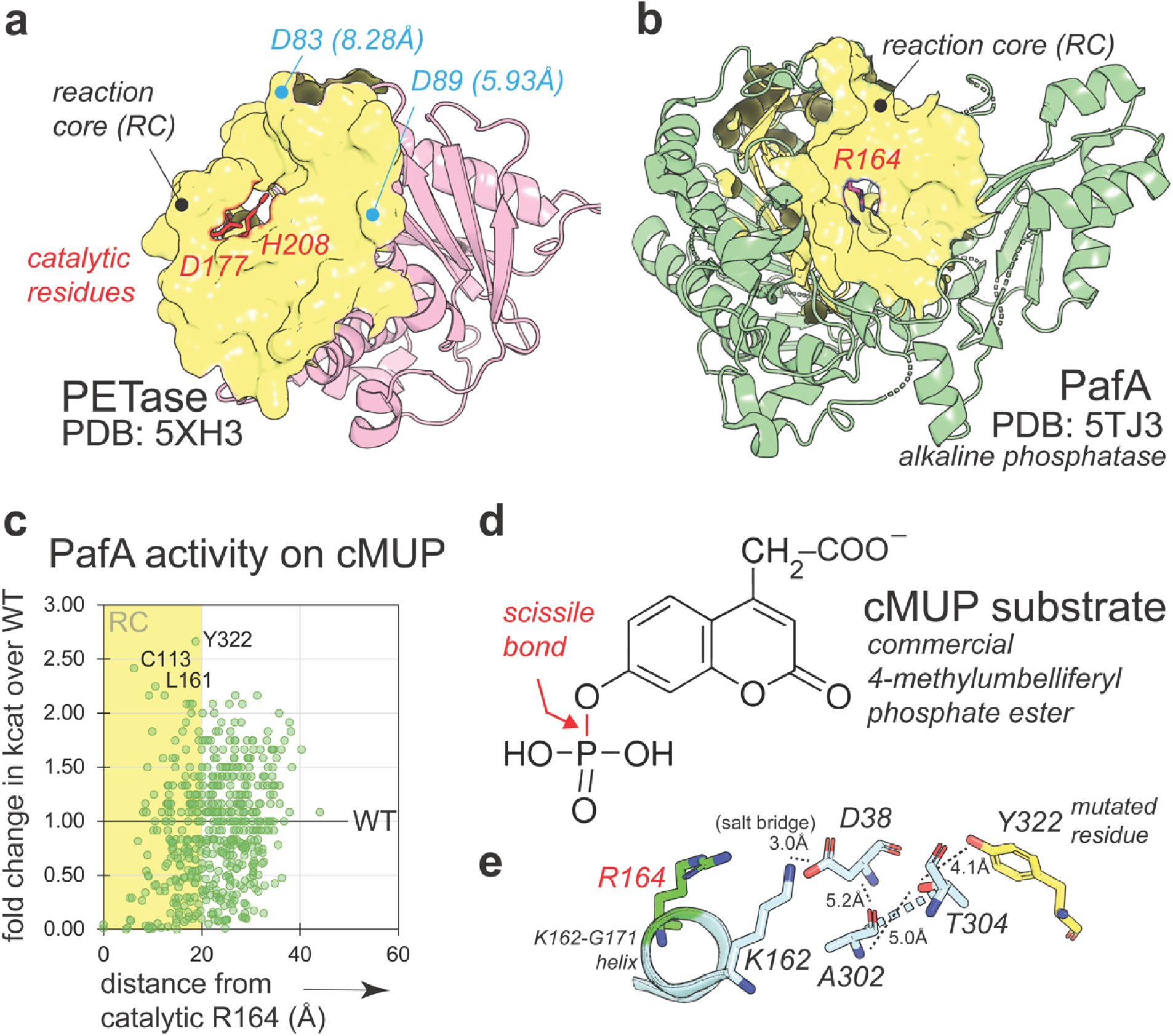
Justification of appropriateness in reaction core definition to capture distal catalysis-enhancing mutational loci. **a)** Reaction core of the PET hydrolase (PDB: 5XH3), marked as yellow surface. The two catalytic residues – His208 and Asp177 are marked in red. Mutations to Asp83 and Asp89 are positions known to improve experimental turnover although >5.8Å away from the active site. We show that our definition of reactive core (RC) for this enzyme would successfully capture both these catalytically beneficial loci despite them being distal to the active site. **b)** RC of industrial alkaline phosphatase is shown with catalytic Arg164 marked in red. **c)** Variation of maximum experimental fold change in cMUP-breaking activity from mutations to all 522 enzyme residues placed at different distances from the catalytic Arg164. The yellow region indicates residue positions within the reactive core. Tyr322, Cys113, and Leu161 mutations show highest increase in catalytic efficiencies. **d)** Structure of the cMUP substrate which is cleaved by PafA at the phosphate bond (shown in red). **e)** Non-covalent contact/ allosteric map of electrostatic forces which provides a structural underpinning of how Tyr322 mutations are likely to impact catalytic activity at Arg164 from a ∼20Å distance.

Next, we investigated the utility of RC on alkaline phosphatase (PafA) from *Elizabethkingia meningoseptica* expressed in *E. coli.* 1039 variants across all 522 residue positions in PafA were experimentally assessed for enzyme activity in cleaving a commercial 4-methylumbelliferyl phosphate ester^45^. It is interesting to note that the top five mutants (in terms of fold increase in activity over wildtype) are all >6.25 Å away from the catalytic residue (i.e., outside the active site) but are contained within the RC (**Figure 2**). Among all the twelve mutational loci which led to ∼2-fold increase in enzyme activity, five of them had were outside the RC. However, all their non-covalent (allosteric) linkage networks to the catalytic Asp164 pass through the allosteric paths for the top five mutants all of which are within the RC. For instance, the allosteric path for Tyr322 (top mutational locus) connects Thr304, followed by Ala302 and Asp38 backbone-backbone electrostatic interaction between parallel beta sheets (**Figure 2**). The Asp38 is salt-bridged with Lys162 which consequently anchors the 10 amino acid long, K162-G171 helix (KG-helix). Dynamic positioning of this KG-helix is crucial for appropriate orientation of catalytic Asp164 and consequently catalytic efficiencies. Comparison of experimental catalytic efficiencies of PafA against two additional substrates^45^ (see **Supplementary Figure 1**) also corroborate that the top performing mutated residue positions are either always within the RC, or principally coordinate activity through such key residues.

### Statistics of the RC-Hydrolase dataset

In total 40,196 hydrolases have been reported till date (July 2024) in human, bacteria, and archaea. Hydrolase 3.4s (proteases) happen to be the most popular member among human hydrolases (41%), which probably alludes to the research focus on proteases for digestive, immune-therapeutic, tissue engineering efforts that prompted researchers to crystallize human proteases ^46^. Similarly in the bacterial world, proteases (EC 3.4) being are largely studied for evaluating degradation of antimicrobial peptides. While there are additional seven Hydrolase functional EC sub-classes (3.7 through 3.13) including breakers of carbon-carbon bonds (EC3.7), sulfur-nitrogen bonds (EC3.10), carbon-sulfur bonds (EC3.13) and so on, most of them (>57%; 23415 instances) belong to 3.1-3.6 sub-EC classes. These structures are from nineteen different organisms, with the majority (48%) from *Homo sapiens*. Host distribution of hydrolases is enumerated in **Figure 3**.

**Figure 3.**
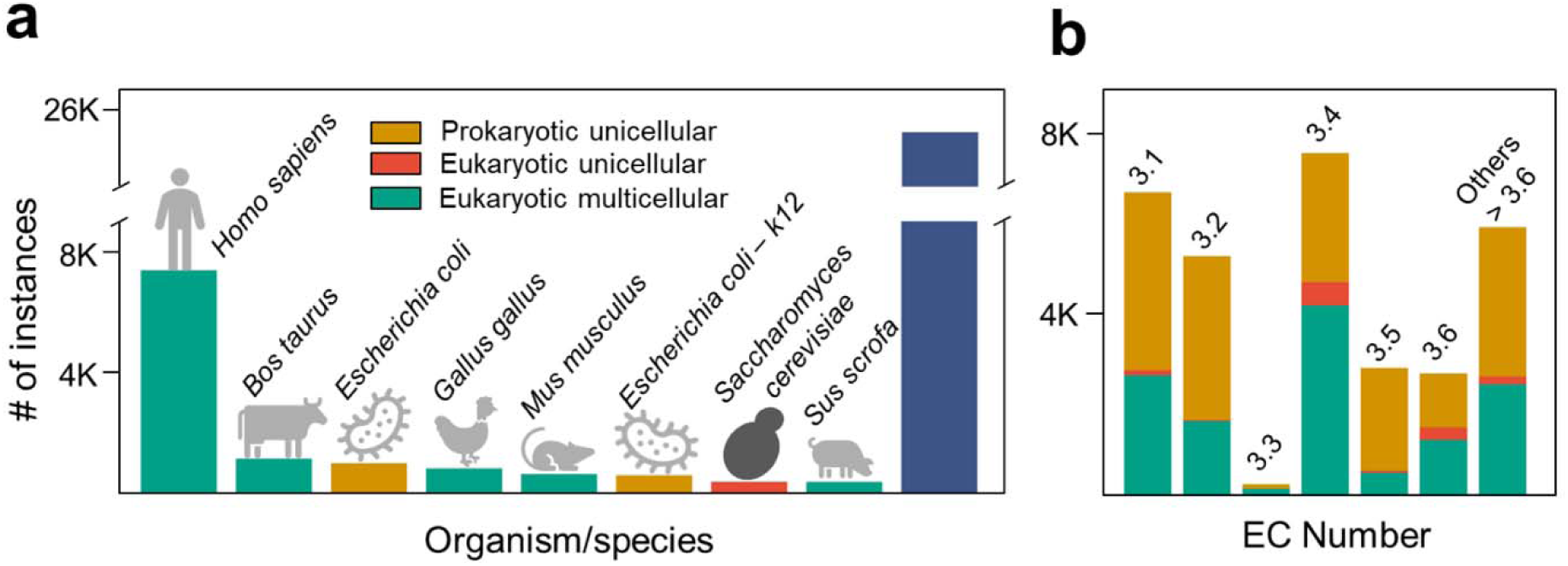
Data statistics of the hydrolase enzymes with known experimental structures. **a)** Demography of structures across eight organisms relevant for synthetic biology and biomedical applications. The blue broken bar at right indicates all other organisms **b)** Number of hydrolases across all thirteen sub-EC classes with host-specific distributions for 3.1 – 3.6 and others.

### Pairwise reactive core similarity across hydrolases from different organisms and EC classes

We centered this study around the preliminary hypothesis that – the function and performance of an enzyme is driven by the shape of the reaction core (catalytic groove and the neighboring allosteric shells spanning 20Å). Thus, hydrolases belonging to the same sub-EC class acting on similar substrate are likely to have similar RC structure while different EC-class hydrolases significantly different RC structure. An inter-EC class RC match between two differently functioning RCs are therefore rational starting anchors to design bifunctional enzymes. Consequently, pairwise similarity scores between all RC structures provides us with insight into potential versatile, engineerable enzyme backbones. All against all matching of 16662 (isoform removed) instances yields 128,568,630 pairwise comparisons. A cut-off of 60% similarity (RCSim >0.6) score between RC structures as per our CADSEEK method, results in 155,329 pairs. This shows than only 0.12% of total 128M enzyme pairs have high RC similarity for mutual engineerability of function. This indicates (a) high diversity catalyzed substrate chemistry results in low similarity of the reaction core (RC) shapes overall, but (b) a tractable, machine learning-worthy list of natural enzymes to query for multifunctional enzyme design.

High RC similarity within the same EC class dominate the pair list (93% cases), as shown by blue scores reported along the diagonal of the heatmap (**Figure 4**). Nearly all (>85%) pairwise comparisons reveal a RC similarity of 60-70%. We divided all ligand and host organism annotations of these hydrolases into additional classes (see https://rc-hydrolase.onrender.com/) relevant for substrate-specificity engineering applications of natural enzymes. There were ∼2.5% pairwise instances (3,889 cases) where two hydrolases of different function (EC number) had more than 70% similarity in reactive core structures. Despite this being an overall small subset, these represent the highly engineerable cases where not only switching host is possible, but also a complete switch of function is likely. Owing to the high similarity in their overall catalytic groove and surrounding allosteric shells (reactive core), we pinpoint these to be amenable for designing promiscuous ^47^ or multifunctional enzymes. This represents the first systematic, structure-guided annotation of natural, promiscuous enzymes. In this respect, the esterase class EC3.1 is most diverse **(Figure 4),** having highest similarity with the other five EC classes. A possible reason might be the broad definition of the EC class 3.1 consisting of nucleases, phosphatases and lipases thus resulting similarity in shape with other EC classes thereby implicitly representing a wide portfolio of acceptable substrate sizes, chemistries, and conformations.

**Figure 4.**
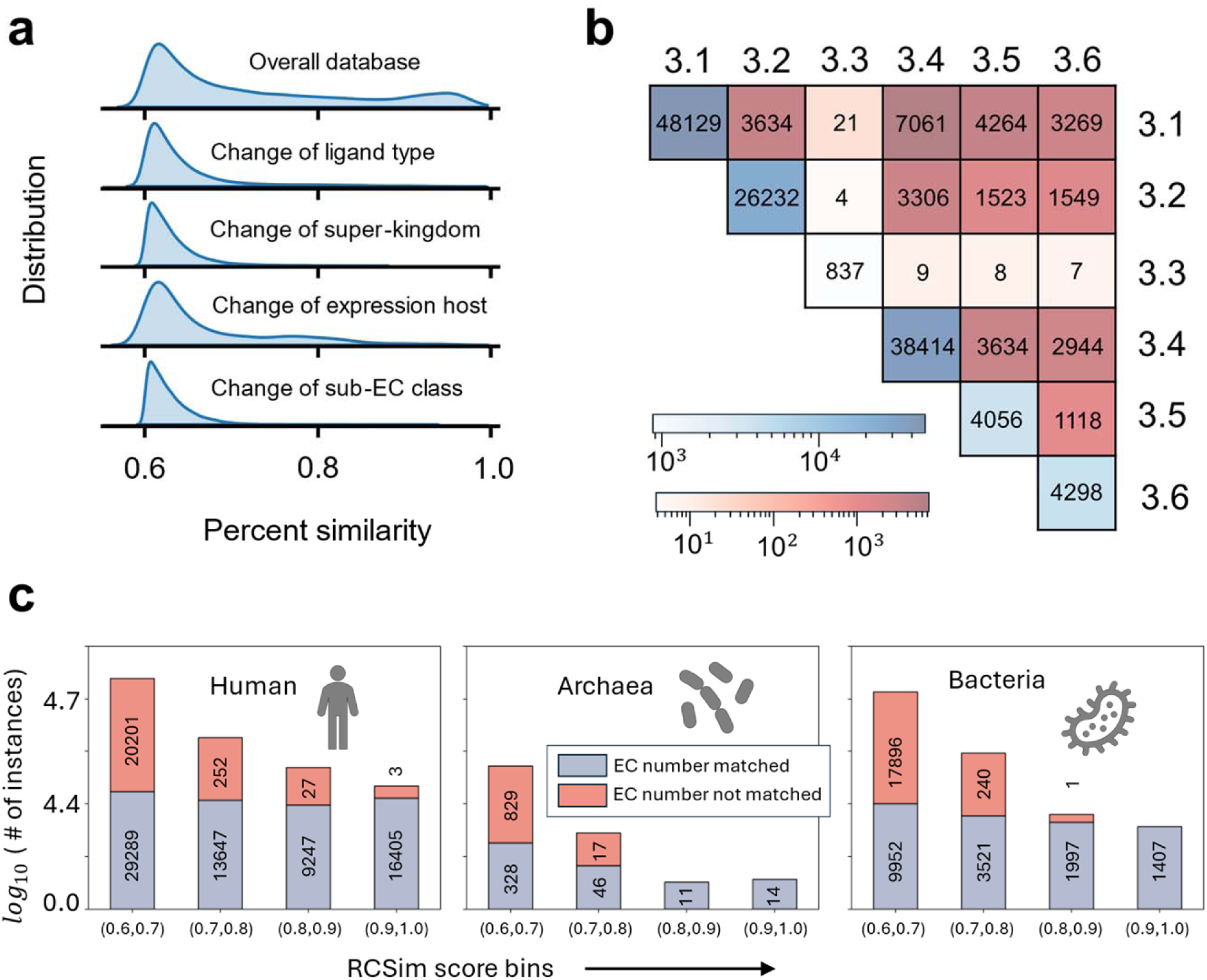
Distribution of percent similarity of pairwise reaction core shapes in the database after 60% RCSim score cutoff across all known hydrolase pertaining to the six major EC classes 3.1-3.6. **a)** Number of hydrolase pairs with known structures with varying (>60%) structural similarities of RCs across different ligand types, expression host organisms, four main super kingdoms, and substrate chemistries. **b)** Number of hydrolases which share more than 60% RCSim score within or across functional EC classes are represented in the heatmap. The diagonal elements show comparison within members of same EC class (blue log scale) and the off-diagonals show hydrolase pairs which show high RC similarity across different EC-classes (red log scale). **c)** Hydrolase pairs expressed within human, archaea, and bacteria with similar (gray) and different (red) functions arranged in RC similarity bins (60% through 90%) with 10% increment.

### Hydrolases with similar reaction cores structures but different expression hosts

Choice of an appropriate enzyme expression host is crucial in synthetic biology owing to their impact on production yield, fermentation scalability, efficiency, and protein functionality^48^. High-yielding hosts like *E. coli* are cost-effective and rapid but may lack necessary post-translational apparatus such as glycosylation, often essential for enzyme functionality thus needing mammalian expression systems (CHO cells^49^). The choice of host can also affect activity levels of enzymes, with some hosts better facilitating appropriate folding and disulfide bond formation^50^. Cost and scalability are also considerations, with bacterial/ fungal systems generally cheaper and faster than mammalian systems. Additionally, host compatibility with the enzyme of interest is important, as production of some enzymes may be uncoupled to growth in some hosts thus requiring either strain design or host switching to ensure acceptable production titers ^48^. Regulatory approval also favors well-established hosts such as CHO cells for therapeutic proteins^51^ or *Bacillus subtilis* for detergent enzymes^52^. The ease of protein purification can also vary, with hosts like yeast and bacteria (e.g., *Pichia pastoris*, Sf9 cells using *Baculovirus*) providing a balance between cost and the ability to perform complex modifications^53^. Selecting the right host-enzyme combination ensures acceptable production titers of a given protein with the desired activity, impacting the success of therapeutic and industrial research end goals.

Having a query-able database that clusters similar reactive core structures across multiple expression hosts is therefore key to pinpointing starting, functional enzyme(s) sequences which are more likely to have appropriate folding and post-translational modifications. For instance, if a target enzymatic conversion requires specific glycosylation patterns on the enzyme, RC-Hydrolase database can be used to find mammalian homologs of a known functional hydrolase. This approach thus allows for the selection of hosts across multiple super-kingdoms of life, *viz* prokaryotes or eukaryote to streamline industrial synthetic biology. As an exemplar, using RC-Hydrolase database, we have identified a pair of ATPase enzymes from two different expression hosts *M. smegmatis* (prokaryotic; PDB accession 6FOC) and *S. cerevisiae* (eukaryotic; PDB accession 2HLD). It is particularly noteworthy that these two enzymes share low overall sequence (60.72%) and structure (RMSD >8Å, GDT_TS 0.29) similarity and would not be classified as any current enzyme comparison metric to be similar besides EC number. However, these two enzymes show extremely high RCSim score (77.9%; 99.9^th^ percentile) indicating very high structural similarity of reactive core (**Figure 5**).

**Figure 5.**
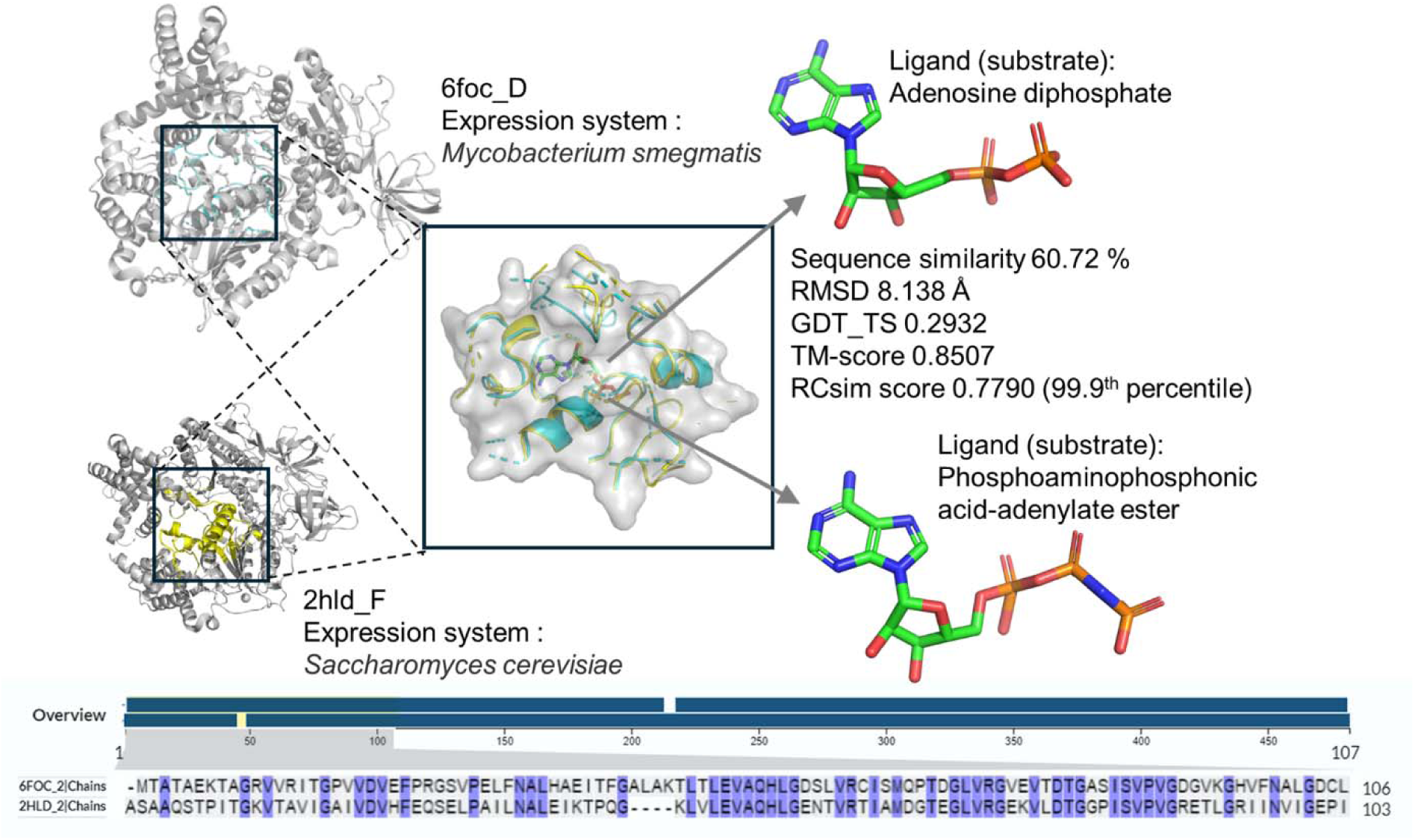
Two protein structures from different expression hosts - *M. smegmatis* and *S. cerevisiae* have structurally similar reaction core with RCSim score >0.77 (99.9^th^ percentile) even though their overall low structural similarity (85%-fold similarity, only 30% global distance match, and >8Å dissimilarity in atomic coordinates) and low sequence similarity (∼60%). Such pairs would be categorized as dissimilar by every standard metric despite having a very similar reactive core structure. RCSim score can thus be leveraged to enable eukaryote-prokaryote host switching in industrial production settings.

### Computationally designing more than one function in a synthetic enzyme and its utility

From the curated RC-Hydrolase database we take a pair of hydrolase enzymes which have different sub-EC classes (i.e., functions) -MTH1 phosphatase (EC3.6; PDB accession 6JVN) and methionine aminopeptidase (MetAP) (EC3.4; PDB accession 5D6E) both originally expressed in *Homo sapiens* (**Figure 6**). Although they have very low sequence similarity (19.6%), and low overall structural similarity (low GDT_TS score 0.0795, high RMSD ∼6.9Å), their reaction core structures have a very high shape similarity (RCSim score 0.7164, 99.5^th^ percentile across all 155,231 hydrolase pairs). Owing to very different substrate choices, their catalytic apparatus is also different albeit housed in a similar looking pocket shape. MTH1 phosphatase is one of the “housekeeping” enzymes that hydrolyze damaged nucleotides in cells and thus prevent them from being incorporated into DNA^54^. MTH1 is a major cancer biomarker and inhibition of MTH1 has wide applicability in the treatment of cancer^55^. MTH1 phosphatase (6JVN) uses a Tyr7 for catalysis. MetAP (5D6E), on the other hand is responsible for the co-translational cleavage of initiator methionines from nascent proteins^56^. The MetAP subtype-2 is up-regulated in many cancers, and selective inhibition of MetAP2 suppresses both vascularization and tumor growth^56^. MetAP uses Asp262 and His331 as the catalytic dyad. Despite lacking sequence-structure homology, both enzymes share a high RCSim score indicating very similar reaction core shapes. Therefore, they serve as ideal design anchors to candidates for accommodating two types of substrates (nucleotides and methionine termini of peptides). Given the high RC similarity, they could serve two different design goals - (a) small molecule inhibitor design with therapeutic utility in arresting cancer, and (b) bi-functional enzyme design with biotechnological utility in removing nucleotide and peptide impurities in industrial carbohydrate production pipelines.

**Figure 6.**
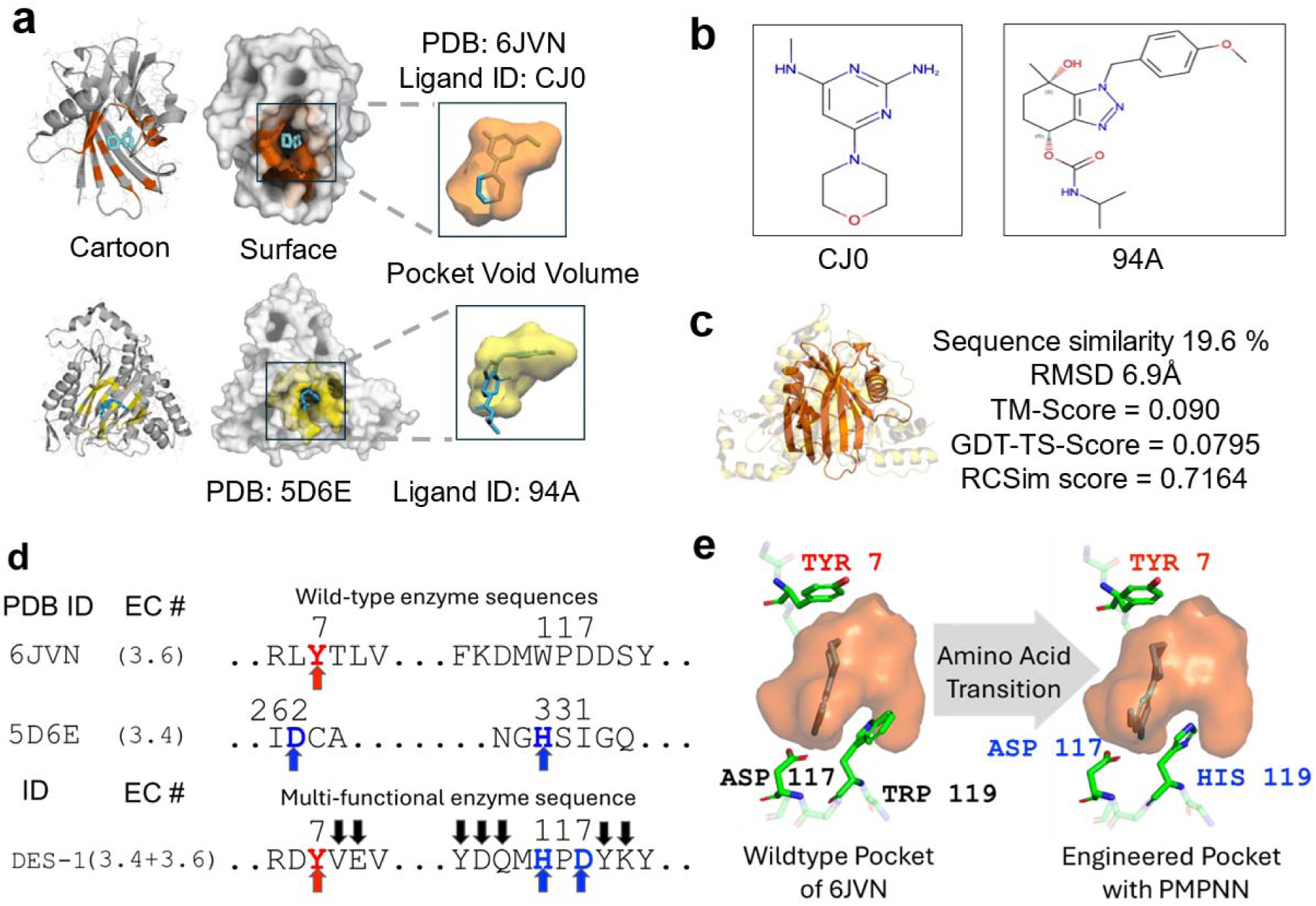
Design pipeline proposition for novel engineered enzyme using RC-Hydrolase database and ProteinMPNN. **a)** Two hydrolase structures - 6JVN (EC3.6 MTH1 nuclease) and 5D6E (EC3.4 MetAP peptidase) share similarity in their reaction core shape and the reaction core void volume. **b)** Chemical structure of the respective inhibitor ligands-CJ0 for 6JVN and 94A for 5D6E. **c)** There is little similarity between the two proteins in terms of sequence and structure as per sequence similarity, RMSD score, TM-score and GDT-TS score. The structure of reaction core is similar as shown by RCSim score 0.7164 which is in 99^th^ percentile of similarity across all hydrolase with known structure. **d)** Reaction core for MTH1 sequence shows catalytic residue Tyr7 marked with red, and MetAp catalytic residues Asp262 and His331 marked with blue arrows. The most stable computationally designed *de novo* bifunctional RC shows both catalytic motifs houses antipodally.

For the first design goal, knowledge that MTH1 shares a high reaction core similarity with MetAP, can be leveraged to design biologics-based inhibitors for either case. Using native substrate structure and conformational isomers of MTH1 one could envision chemo-inspired inhibitors design for MetAP using conditional deep generative models like PMDM ^57^ and DrugDiff^58^. Similarly, MetAP substrate information can be now used to design similar inhibitors for MTH1. Given the pressing need for potent cancer inhibitors, RC-Hydrolase can be used as a cornerstone facilitate expedited inhibitor discovery that target multiple cancer pathways from a single structural analysis.

For the second, more ambitious design goal, we envision the MTH1-MetAp RC pair can be leveraged to *de novo* design promiscuous enzymes that can cleave both nucleotides and methionines from peptides. Prior sporadic success on experimental design of such multifunctional enzymes indicates that there must be additional scaffold engineering to sustain the *de novo* reaction core^59^. Notwithstanding, we now provide a structurally guided protocol to design the multifunctional reactive core.

Residue-by-residue matching at the RC groove lining between MTH1 and MetAP reveals that the Trp119 in MTH1 (6JVN) sequence when altered to His119 results in synergistic rotamer repacking with neighboring native residue Asp117 to identically mimic the side chain dihedrals of catalytic dyad His331 and Asp262 of wildtype MetAP. This MetAP catalytic dyad is stably (as per AlphaFold2) housed within the RC of MTH1 and positioned antipodally to MTH1’s own catalytic Tyr7 (**Figure 6**). This structurally-crafted *de novo* reactive core is further stabilized using ProteinMPNN to through additional compensatory mutations to arrive at a library of nuclease-protease bifunctional MTH1 variants.

ProteinMPNN was used to design an ensemble of sequences that can adopt the Trp119His amino acid transformation in the crystal structure of MTH1. ProteinMPNN was used to only mutate RC residues while keeping the three catalytic residues (Try7 for native MTH1 EC3.6 nuclease activity, and Asp262 and His119 for EC3.4 MetAP peptidase functionality). The surface-exposed MTH1 scaffold outside the RC was kept unaltered for this exemplar. Using a sampling temperature of 0.3, we generated 65 novel sequences (see **Supplementary table 1**). One of the fifteen sequences showed high stability (ProteinMPNN global score >1, average pLDDT > 0.87, RMSD <0.9Å with native MTH1).

### RC-Hydrolase database description

RC-Hydrolase is the first-of-its kind publicly available comprehensive enzyme database (available : https://rc-hydrolase.onrender.com) which contains reaction core annotations and RC sub-structures. It features all hydrolases spanning sub-EC class 3.1 through 3.6 (**Figure 7**) with experimental confirmed crystal structures. Here reaction cores of hydrolases are clustered as per their explicit structural similarity with an implicit knowledge of the chemical groups that line these reaction cores. CADSEEK instantaneous pairwise shape match engine enables rapid search and retrieval of identical or similar reaction core geometries.

**Figure 7.**
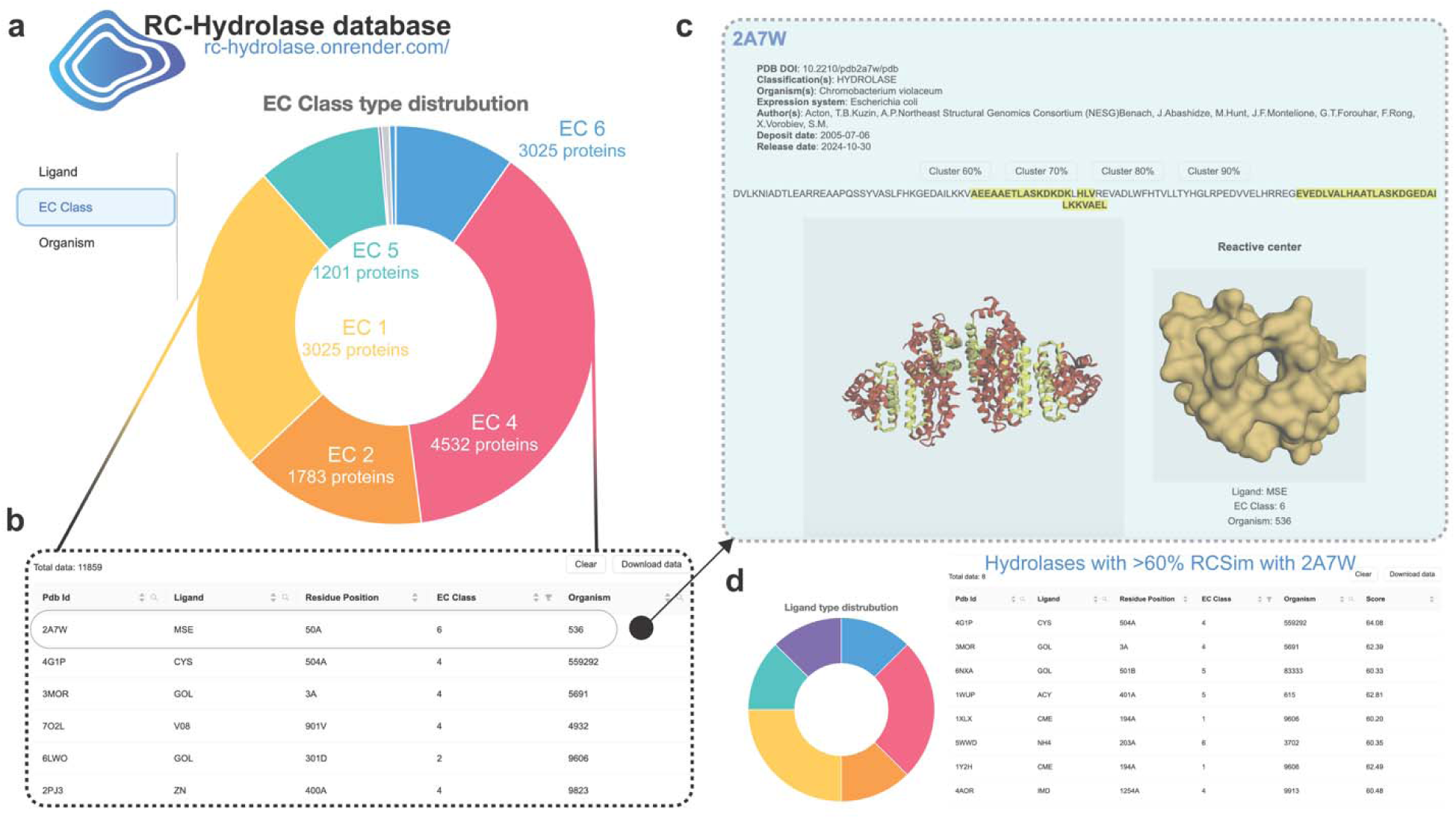
Snippet from the RC-Hydrolase database. **a)** A graphical overview of number of reaction cores of hydrolases when clustered by their ligand, EC Class, and expression host organism information. b) A tabulated, downloadable version of the data with reported PDB accession ids, bound ligand, InterPro-annotated catalytic/ active site residue, EC class, and expression organism id (for example, 9606 is *Homo sapiens*) is listed. **c)** PDB metadata about source research article, organism, expression system, sequence, and overall structure of the enzyme is listed. The reactive core is highlighted in yellow, and the RC sub-surface is shown on the right inset. **d)** Clustering a sample hydrolase with PDB accession 2A7W all other hydrolase reactive cores with at least 60% RC similarity reveals eight natural enzymes. Their EC numbers, expression hosts, and RC Sim scores are reported. Sequences, overall structures, and RC substructures of this cluster are downloadable from this page.

## Conclusion

Hydrolases play a crucial role in industrial applications, making up nearly 75% of all enzymes used in industry. Among these, amylase, proteases, and lipases are the most prevalent, together representing more than 70% of the enzyme market^60^. Their widespread use underscores their importance in various industrial processes. Numerous techniques have emerged for experimentally or computationally determining whole protein structures from sequences. However, a database of reaction cores of biocatalytic proteins does not exist yet. This has forced data-driven, enzyme activity prediction to be sequence-driven only (for example, DLKcat ^61^). CatPred^62^ attempts to incorporate whole enzyme structures for activity prediction and demonstrates a loss of prediction accuracy when compared to sequence-only approaches. This work puts forward a comprehensive database of structural folds of catalytically relevant sub-structure from all experimentally reported hydrolase structures till date. Reasoning over such focused sub-structures (reactive cores - RCs) has been shown to be highly informative in better functional clustering of enzymes in this study. We also laid out a putative computational recipe to effectively design small molecule inhibitors and *de novo* promiscuous (multifunctional) enzyme reaction cores. RC-Hydrolase database allows users to down-select easy-to-engineer natural hydrolase backbones. Careful selection and design of such hydrolase backbones from reliable industrial expression host organisms can open novel frontiers for successful design of highly active multi-functional hydrolase reactive core folds. Such designed multi-functional reactive cores can be further stabilized using protein hallucinators (ProteinMPNN). This also extends onto taking stable non-enzymatic backbones and endowing them with catalytic motifs (such as design of catalytic antibodies). In future we intend to expand our RC-Hydrolase database to include reaction cores of AlphaFold2-predicted hydrolases (4.33M additional entries as per UniProt).

In the biotechnological industry, single enzymes with multiple functions have garnered significant interest due to their versatility and efficiency ^63^. These multifunctional enzymes are designed by grafting different hydrolase catalytic sites onto one enzyme molecule, enabling them to catalyze multiple reactions promiscuously. Final stable variants are currently identified only through heuristics or directed evolution. Promiscuous enzymes which pack in more than one function reduces the need for multiple enzymes and streamlines the purification steps for commercial bioproducts (proteins, lipids, carbohydrates, and nucleotides). For instance, a single enzyme engineered to possess both protease and lipase activities, allows it to break down both proteins and fats. This capability is particularly valuable in industries such as food processing, detergent manufacturing, removal of recombinant DNA from protein products, removal of impurities from drugs, and biofuel production, where complex substrates are common. Additionally, these enzymes can be tailored to have condition-specific activities (distinct pH windows for differential functional identities), enhancing their versatility for industrial applications. We hereby compile a crucial database to enable future advanced enzyme design campaigns. We will continually update RC-Hydrolase database with new data from PDB, non-hydrolase enzymes (such as oxidoreductases and transferases), and even include non-catalytic grooves from enzymes which bear high similarity to known RCs. The entire enzyme will thus become “free real estate” for *de novo* generation of catalytic sites. Strides in structural bioinformatics will include topology matching, overlaid on biochemical information rather than homology-based sequence classification. As the field of generative AI based protein engineering grows, we hope to use the RC-Hydrolase database to design highly promiscuous and microenvironment-driven, conditionally active reaction cores. .

In immediate future, RC-Hydrolase will require synergistic experiments to inform the minimum RC similarity (naive threshold set at 60%) to warrant engineerability. We seek to fine tune the geometric features that impact the reaction such as, cut-off radius for reaction core to see how it impacts the pairwise comparison (RCSim scores). We focused on ligands for catalytic relevance, with metals cofactors and small molecule inhibitors. In future, detailed simulations that explore the energetic contribution of water molecules, towards hydrolysis would be accounted *via* density functional theory measurements, towards the RCSim score.

## Methods

### Extracting reaction cores from whole enzyme structures

The first goal of this work is to identify a known, conserved domain or substrate binding site in the protein structure. Conserved domains in protein sequences are annotated and can be accessible through several databases but are not mapped onto protein structures. Each PDB file often contains several chains of protein along with heteroatoms. Conserved domain and hetero atom locations are taken as guides to carve out regions of the protein surface as the catalytic core. The specific steps are outlined as follows.

*<u>Step 1.</u>* For each chain in every hydrolase entry in the Protein Data Bank, the sequence of each chain is extracted and conserved domains in the chains are cross annotated using Interpro enzyme catalytic database. If catalytic annotation is not found for any entry, a family sequence alignment is first performed to find the closest homolog which has an Interpro annotation to pinpoint the catalytic residue for the query enzyme.

*<u>Step 2.</u>* Enzymes are structurally and functionally very diverse. We aimed identify the region across all structural fold types, which is closest to the ‘center of activity’. Beyond conserved catalytic residues, co-crystallized heteroatoms have been used as structural cues in locating the site of activity. Because heteroatoms other than water mostly comprise metal ions which serve as enzyme co-factor or substrate analogs/ inhibitors both of which preferentially bind to the active site. Associated challenges include active site grooves that are created at the interface of multimeric enzyme chains. Hence one cannot simply assume this process to be single chain basis.

*<u>Step 3.</u>* From a bound, annotated molecule or the confirmed catalytic residues a search radius of 20Å is taken and the number of conserved residues in that range is counted. In case of multiple bound small molecules at different pockets on an enzyme structure, we have empirically established that the pocket with greatest number of conserved residues (from family sequence alignments) within 20Å is always the active site.

*<u>Step 3.</u>* After the principal groove is identified, all residues that are connected to its neighbors *via* electrostatic interaction or hydrophobic packing within a 20Å diameter are extracted and defined as the reaction core (RC). The radius is selected by an educated guess taking into consideration that no major contributor to the active site fold is missed. A hard fraction of the total number of residues in the chain or the total structure also is not a great choice, as there are protein instances where the whole chain acts as a conserved domain and also where a domain contains only one residue from a chain.

### RC-Hydrolase database

We started with the PDB database. As of 1^st^ July 2024, there were 40,196 instances of known hydrolase structure on three important spans of life – archaea, eukaryote, and bacteria, with 26,143 single chain, 14,028 multi-chain hydrolases spanning short peptides (six amino acid long) to multi-chain complexes (4426 amino acids). Among these there were minimum instances where structure had several uncrystallized/ missing residues, and even corrupted structure files - which were all removed. Overall catalytic mechanisms include – nucleophiles with a defined catalytic triad (serine protease, cysteine protease etc.), activating water molecule with coordinated metal ions (matrix metalloproteinases/ MMPs). However, the lack of a curated Hydrolase reaction core (RC) structure database prevented reliable functional clustering and latent structural encoding of enzymes for function prediction.

### CADSEEK 3D shape search and analytics engine for automated encoding and classification of RC-Hydrolase database

CADSEEK is a highly efficient and scalable 3D shape search engine capable of processing any type of 3D digitized shape. It extracts a shape code from the digitized geometry and automatically builds a classification without AI training. This enables users to search for identical or similar 3D shapes in under 2 milliseconds, optimizing the utilization and mining of 3D digital assets. The algorithm mathematically codifies a 3D shape by converting it into a boundary value problem. The solution function of the boundary value problem is expressed in series form, where the coefficients of the series expansion are unique representatives of the boundary conditions. The 3D shape surfaces are used as the boundary conditions for the series expansion solution where an electrostatic charge distribution over the 3D object surfaces simulates the boundary conditions. The field satisfies Poisson’s equation for which the solution is unique and non-invertible for a given boundary condition ^64^. Extracted shape codes are then automatically classified by CADSEEK’s algorithms to generate a search index.

RC-Hydrolase dataset consisting of >16k 3D protein reaction core shapes. Analysis time on a on a single Quad core i7 or similar was as follows: extracting RC-Hydrolase 3D shape codes took 4 compute hours, the automated classification to generate a searchable index required 10 compute minutes, and the 3D shape analytics report for the full dataset was generated in under 4 seconds. CADSEEK shape coding can be threaded and can be parallelized across multiple workstations or computers.

## Supplementary Data Statement

Supplementary Data is available. PDB file of the extracted pockets and data sheet of result can be accessed in the supplementary files.

## Data Availability

The database can be accessed at - https://rc-hydrolase.onrender.com/. This has been also listed in the PDB under *Structure Classification Analyses* within *Additional Resources*.

## Declaration of generative AI and AI-assisted technologies in the writing process

During the preparation of this work the author(s) used no generative AI tools and take(s) full responsibility for the content of the publication.

## Author Contributions

R.C. conceived and designed the study. S.F. and P.G co-developed all scripts. K.A.S. helped in annotating the reaction centers and provided critical advice on *in silico* design strategy for the multifunctional enzymes. V.S.R. performed the analysis on industrial PafA enzyme. K.H. helped with downloading and analyzing the PDB structures. Y.C.T made the latest rendition (rc-hydrolase.onrender.com) RC-Hydrolase database. A.Q. performed the automated 3D reaction center shape encoding, automated classification, and generating shape analytics report to perform 3D pocket shape comparisons and set up the RC-Hydrolase dataset on iSEEK’s 3DshapeIndex.com. N.F.R., N.A., and M.K. provided key insights about experimental insights to the study. All authors wrote and edited the manuscript drafts and agree with findings reported in the manuscript.

