## Supplemental File 1 for "Improved Functional Classification of Hydrolases through Pairwise Structural Similarity of Reaction Cores"

^4^iSEEK Corporation, Ames, IA, USA

^5^Department of Mechanical Engineering, University of Texas at Austin, Texas, USA

^#^ These authors contributed equally

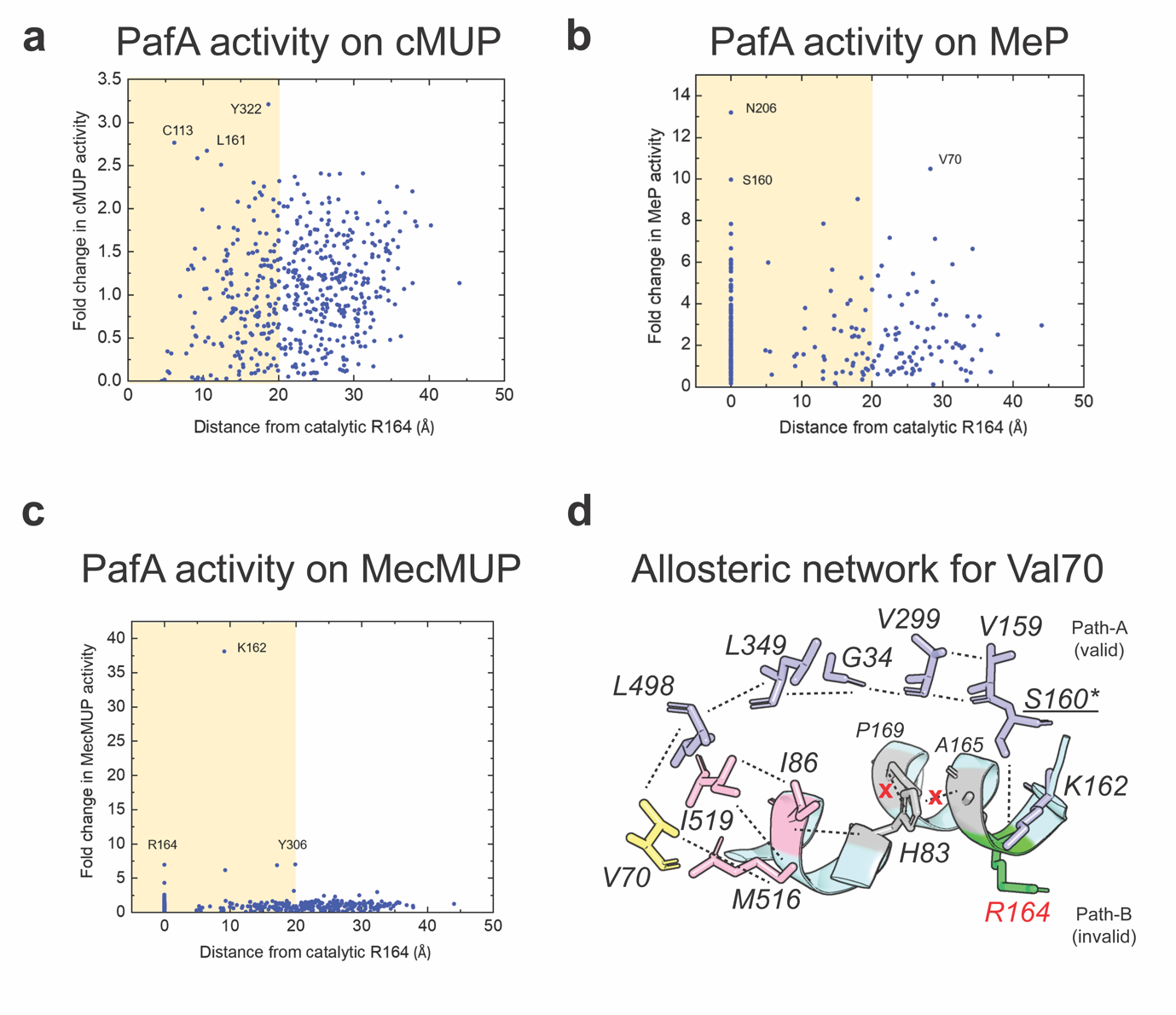
**Figure S1** Variation plots of maximum experimental fold change in substrate-braking activity of different mutant residues, each at a different distance from the catalytic residue Arg164 (R164) corresponding to hydrolysis of three different substrates **a)** cMUP: Carboxy 4-methylumbelliferyl phosphate ester- a binding limited substrate, **b)** MeP: methyl phosphate- a chemistry limited substrate and **c)** MecMUP: Methyl Phosphodiester. The yellow region indicates residue positions within the reactive core (RC). **d)** Allosteric network analyses for V70 shows that there are two possible routes for V70 to interact with catalytic R164. In the first route (Path-A) a series of anti-parallel beta pleats provide a stable backbone-mediated hydrogen-bonded path to V159. This path is further strengthened by side-chain hydrophobic packing between all affiliated residue positions. V159 is covalently linked to S160 (which is within the RC and is a strong regulator of catalytic efficiency). S160 is in turn electrostatically linked to K162 which is part of the helix that contains the catalytic R164. In addition, the Path-B shows a weaker side-chain-mediated allosteric path up to H83. However, histidine pose no affinity for its neighboring P169 or A165 residues (except for van der Waal's clashes). This explains why V70 position shows only marginally higher catalytic rates over S160. Albeit, the enhanced catalytic ability (with respect to S160) is explained by the additional hydrophobic packing in Path-A and van der Waal's contribution from Path-B.

**Table S1** Protein MPNN-generated 65 sequences of the predicted bifunctional enzyme which folds into the desired pocket shape with two antipodally placed catalytic motifs (one for each function).

| **Fastaheader** | **Sequence** |
| --- | --- |
| 6jvn-W116H, score=1.5857, global_score=1.4311, fixed_chains=[], designed_chains=['A'], model_name=v_48_020, git_hash=8907e6671bfbfc92303b5f79c4b5e6ce47cdef57, seed=37 | ASRLYTLVLVLQPQRVLLGMKKRGFGAGRWNGFGGKVQEGETIEDGARRELQEESGLTVDALHKVGQIVFEFVGEPELMDVHVFCTDSIQGTPVESDEMRPCWFQLDQIPFKDMHPDDSYWFPLLLQKKKFHGYFKFQGQDTILDYTLREVDTV |
| T=0.3, sample=1, score=0.7570, global_score=1.3587, seq_recovery=0.6154 | ASRLYTLVFCLQPQRVLLGMKKRGFGAGRWNGFGGKVQEGETIEDGARRELQEESGLTVDALHKVGQIVFEFVGEPELMDVHVFCTDSIQGTPVESDEMRPCWFQLDQIPFKDMFPDFSLWFPLLLQKKKFHGYFKFQGQDTILDYTLREVDTV |
| T=0.3, sample=2, score=0.8535, global_score=1.3392, seq_recovery=0.5385 | ASRLYTLVFCLQPQRVLLGMKKRGFGAGRWNGFGGKVQEGETIEDGARRELQEESGLTVDALHKVGQIVFEFVGEPELMDVHVFCTDSIQGTPVESDEMRPCWFQLDQIPFKEMFPDFSKWFPLLLQKKKFHGYFKFQGQDTILDYTLREVDTV |
| T=0.3, sample=3, score=0.7372, global_score=1.3424, seq_recovery=0.6154 | ASRLYTLVFCLQPQRVLLGMKKRGFGAGRWNGFGGKVQEGETIEDGARRELQEESGLTVDALHKVGQIVFEFVGEPELMDVHVFCTDSIQGTPVESDEMRPCWFQLDQIPFKQMHPDFSLWFPLLLQKKKFHGYFKFQGQDTILDYTLREVDTV |
| T=0.3, sample=4, score=0.8718, global_score=1.3840, seq_recovery=0.4615 | ASRLYVLVFCLQPQRVLLGMKKRGFGAGRWNGFGGKVQEGETIEDGARRELQEESGLTVDALHKVGQIVFEFVGEPELMDVHVFCTDSIQGTPVESDEMRPCWFQLDQIPFKQMFPDYSLWFPLLLQKKKFHGYFKFQGQDTILDYTLREVDTV |
| T=0.3, sample=5, score=0.8548, global_score=1.3670, seq_recovery=0.5385 | ASRLYTLVFCLQPQRVLLGMKKRGFGAGRWNGFGGKVQEGETIEDGARRELQEESGLTVDALHKVGQIVFEFVGEPELMDVHVFCTDSIQGTPVESDEMRPCWFQLDQIPFKDMWPDASLWWPLLLQKKKFHGYFKFQGQDTILDYTLREVDTV |
| T=0.3, sample=6, score=0.7136, global_score=1.3390, seq_recovery=0.6154 | ASRLYTLVFCLQPQRVLLGMKKRGFGAGRWNGFGGKVQEGETIEDGARRELQEESGLTVDALHKVGQIVFEFVGEPELMDVHVFCTDSIQGTPVESDEMRPCWFQLDQIPFKDMFPDASLWFPLLLQKKKFHGYFKFQGQDTILDYTLREVDTV |
| T=0.3, sample=7, score=0.6637, global_score=1.3639, seq_recovery=0.5385 | ASRLYTLVFCLQPQRVLLGMKKRGFGAGRWNGFGGKVQEGETIEDGARRELQEESGLTVDALHKVGQIVFEFVGEPELMDVHVFCTDSIQGTPVESDEMRPCWFQLDQIPFKQMWPDASLWFPLLLQKKKFHGYFKFQGQDTILDYTLREVDTV |
| T=0.3, sample=8, score=0.7424, global_score=1.3320, seq_recovery=0.5385 | ASRLYTLVFCLQPQRVLLGMKKRGFGAGRWNGFGGKVQEGETIEDGARRELQEESGLTVDALHKVGQIVFEFVGEPELMDVHVFCTDSIQGTPVESDEMRPCWFQLDQIPFKQMYPDYSLWFPLLLQKKKFHGYFKFQGQDTILDYTLREVDTV |
| T=0.3, sample=9, score=0.8315, global_score=1.3643, seq_recovery=0.6154 | ASRLFTLVFVLQPQRVLLGMKKRGFGAGRWNGFGGKVQEGETIEDGARRELQEESGLTVDALHKVGQIVFEFVGEPELMDVHVFCTDSIQGTPVESDEMRPCWFQLDQIPFKDMYPDASLWFPLLLQKKKFHGYFKFQGQDTILDYTLREVDTV |
| T=0.3, sample=10, score=0.7398, global_score=1.3591, seq_recovery=0.5385 | ASRLFTLVFCLQPQRVLLGMKKRGFGAGRWNGFGGKVQEGETIEDGARRELQEESGLTVDALHKVGQIVFEFVGEPELMDVHVFCTDSIQGTPVESDEMRPCWFQLDQIPFKDMWPDYSLWFPLLLQKKKFHGYFKFQGQDTILDYTLREVDTV |
| T=0.3, sample=12, score=0.7559, global_score=1.3200, seq_recovery=0.4615 | ASRLFTLVFCLQPQRVLLGMKKRGFGAGRWNGFGGKVQEGETIEDGARRELQEESGLTVDALHKVGQIVFEFVGEPELMDVHVFCTDSIQGTPVESDEMRPCWFQLDQIPFKEMYPDYSLWFPLLLQKKKFHGYFKFQGQDTILDYTLREVDTV |
| T=0.3, sample=13, score=0.7302, global_score=1.3677, seq_recovery=0.5385 | ASRLYTLVFCLQPQRVLLGMKKRGFGAGRWNGFGGKVQEGETIEDGARRELQEESGLTVDALHKVGQIVFEFVGEPELMDVHVFCTDSIQGTPVESDEMRPCWFQLDQIPFKNMWPDYSLWFPLLLQKKKFHGYFKFQGQDTILDYTLREVDTV |
| T=0.3, sample=14, score=0.7492, global_score=1.3650, seq_recovery=0.5385 | ASRLYTLVFCLQPQRVLLGMKKRGFGAGRWNGFGGKVQEGETIEDGARRELQEESGLTVDALHKVGQIVFEFVGEPELMDVHVFCTDSIQGTPVESDEMRPCWFQLDQIPFKQMFPDASLWFPLLLQKKKFHGYFKFQGQDTILDYTLREVDTV |
| T=0.3, sample=15, score=0.8550, global_score=1.3907, seq_recovery=0.4615 | ASRLYVLVFCLQPQRVLLGMKKRGFGAGRWNGFGGKVQEGETIEDGARRELQEESGLTVDALHKVGQIVFEFVGEPELMDVHVFCTDSIQGTPVESDEMRPCWFQLDQIPFKEMWPDYSLWFPLLLQKKKFHGYFKFQGQDTILDYTLREVDTV |
| T=0.3, sample=16, score=0.7426, global_score=1.3379, seq_recovery=0.5385 | ASRLYTLVFCLQPQRVLLGMKKRGFGAGRWNGFGGKVQEGETIEDGARRELQEESGLTVDALHKVGQIVFEFVGEPELMDVHVFCTDSIQGTPVESDEMRPCWFQLDQIPFKNMYPDYSLWFPLLLQKKKFHGYFKFQGQDTILDYTLREVDTV |
| T=0.3, sample=20, score=0.7932, global_score=1.3669, seq_recovery=0.4615 | ASRLFTLVFCLQPQRVLLGMKKRGFGAGRWNGFGGKVQEGETIEDGARRELQEESGLTVDALHKVGQIVFEFVGEPELMDVHVFCTDSIQGTPVESDEMRPCWFQLDQIPFKQMFPDFSLWFPLLLQKKKFHGYFKFQGQDTILDYTLREVDTV |
| T=0.3, sample=23, score=0.7867, global_score=1.3665, seq_recovery=0.5385 | ASRLFTLVFVLQPQRVLLGMKKRGFGAGRWNGFGGKVQEGETIEDGARRELQEESGLTVDALHKVGQIVFEFVGEPELMDVHVFCTDSIQGTPVESDEMRPCWFQLDQIPFKQMWPDYSLWFPLLLQKKKFHGYFKFQGQDTILDYTLREVDTV |
| T=0.3, sample=24, score=0.8884, global_score=1.3604, seq_recovery=0.6923 | ASRLYTLVFALQPQRVLLGMKKRGFGAGRWNGFGGKVQEGETIEDGARRELQEESGLTVDALHKVGQIVFEFVGEPELMDVHVFCTDSIQGTPVESDEMRPCWFQLDQIPFKDMHPDYSLWFPLLLQKKKFHGYFKFQGQDTILDYTLREVDTV |
| T=0.3, sample=25, score=0.7106, global_score=1.3470, seq_recovery=0.5385 | ASRLYTLVFCLQPQRVLLGMKKRGFGAGRWNGFGGKVQEGETIEDGARRELQEESGLTVDALHKVGQIVFEFVGEPELMDVHVFCTDSIQGTPVESDEMRPCWFQLDQIPFKNMWPDASLWFPLLLQKKKFHGYFKFQGQDTILDYTLREVDTV |
| T=0.3, sample=26, score=0.7046, global_score=1.3558, seq_recovery=0.6154 | ASRLYTLVFCLQPQRVLLGMKKRGFGAGRWNGFGGKVQEGETIEDGARRELQEESGLTVDALHKVGQIVFEFVGEPELMDVHVFCTDSIQGTPVESDEMRPCWFQLDQIPFKDMWPDYSLWFPLLLQKKKFHGYFKFQGQDTILDYTLREVDTV |
| T=0.3, sample=27, score=0.7411, global_score=1.3585, seq_recovery=0.5385 | ASRLYTLVFCLQPQRVLLGMKKRGFGAGRWNGFGGKVQEGETIEDGARRELQEESGLTVDALHKVGQIVFEFVGEPELMDVHVFCTDSIQGTPVESDEMRPCWFQLDQIPFKQMWPDYSLWFPLLLQKKKFHGYFKFQGQDTILDYTLREVDTV |
| T=0.3, sample=28, score=0.7922, global_score=1.3616, seq_recovery=0.6154 | ASRLFTLVFCLQPQRVLLGMKKRGFGAGRWNGFGGKVQEGETIEDGARRELQEESGLTVDALHKVGQIVFEFVGEPELMDVHVFCTDSIQGTPVESDEMRPCWFQLDQIPFKDMHPDYSLWFPLLLQKKKFHGYFKFQGQDTILDYTLREVDTV |
| T=0.3, sample=29, score=0.8874, global_score=1.3735, seq_recovery=0.5385 | ASRLYTLVFALQPQRVLLGMKKRGFGAGRWNGFGGKVQEGETIEDGARRELQEESGLTVDALHKVGQIVFEFVGEPELMDVHVFCTDSIQGTPVESDEMRPCWFQLDQIPFKEMFPDFSLWFPLLLQKKKFHGYFKFQGQDTILDYTLREVDTV |
| T=0.3, sample=30, score=0.7625, global_score=1.3666, seq_recovery=0.5385 | ASRLYTLVFCLQPQRVLLGMKKRGFGAGRWNGFGGKVQEGETIEDGARRELQEESGLTVDALHKVGQIVFEFVGEPELMDVHVFCTDSIQGTPVESDEMRPCWFQLDQIPFKEMFPDYSLWFPLLLQKKKFHGYFKFQGQDTILDYTLREVDTV |
| T=0.3, sample=31, score=0.7266, global_score=1.3514, seq_recovery=0.5385 | ASRLYTLVFCLQPQRVLLGMKKRGFGAGRWNGFGGKVQEGETIEDGARRELQEESGLTVDALHKVGQIVFEFVGEPELMDVHVFCTDSIQGTPVESDEMRPCWFQLDQIPFKQMYPDFSLWFPLLLQKKKFHGYFKFQGQDTILDYTLREVDTV |
| T=0.3, sample=32, score=0.7560, global_score=1.3671, seq_recovery=0.5385 | ASRLYTLVFCLQPQRVLLGMKKRGFGAGRWNGFGGKVQEGETIEDGARRELQEESGLTVDALHKVGQIVFEFVGEPELMDVHVFCTDSIQGTPVESDEMRPCWFQLDQIPFKEMWPDASLWFPLLLQKKKFHGYFKFQGQDTILDYTLREVDTV |
| T=0.3, sample=33, score=0.7938, global_score=1.3563, seq_recovery=0.5385 | ASRLFTLVFCLQPQRVLLGMKKRGFGAGRWNGFGGKVQEGETIEDGARRELQEESGLTVDALHKVGQIVFEFVGEPELMDVHVFCTDSIQGTPVESDEMRPCWFQLDQIPFKDMYPDFSLWFPLLLQKKKFHGYFKFQGQDTILDYTLREVDTV |
| T=0.3, sample=34, score=0.9521, global_score=1.3634, seq_recovery=0.5385 | ASRLYTLVFCLQPQRVLLGMKKRGFGAGRWNGFGGKVQEGETIEDGARRELQEESGLTVDALHKVGQIVFEFVGEPELMDVHVFCTDSIQGTPVESDEMRPCWFQLDQIPFKQMFPDSSLWFPLLLQKKKFHGYFKFQGQDTILDYTLREVDTV |
| T=0.3, sample=35, score=0.7711, global_score=1.3728, seq_recovery=0.6154 | ASRLYTLVFCLQPQRVLLGMKKRGFGAGRWNGFGGKVQEGETIEDGARRELQEESGLTVDALHKVGQIVFEFVGEPELMDVHVFCTDSIQGTPVESDEMRPCWFQLDQIPFKEMHPDYSLWFPLLLQKKKFHGYFKFQGQDTILDYTLREVDTV |
| T=0.3, sample=36, score=0.7978, global_score=1.3418, seq_recovery=0.6154 | ASRLYTLVFCLQPQRVLLGMKKRGFGAGRWNGFGGKVQEGETIEDGARRELQEESGLTVDALHKVGQIVFEFVGEPELMDVHVFCTDSIQGTPVESDEMRPCWFQLDQIPFKEMYPDYSYWFPLLLQKKKFHGYFKFQGQDTILDYTLREVDTV |
| T=0.3, sample=37, score=0.7518, global_score=1.3413, seq_recovery=0.6154 | ASRLYTLVYCLQPQRVLLGMKKRGFGAGRWNGFGGKVQEGETIEDGARRELQEESGLTVDALHKVGQIVFEFVGEPELMDVHVFCTDSIQGTPVESDEMRPCWFQLDQIPFKDMYPDASLWFPLLLQKKKFHGYFKFQGQDTILDYTLREVDTV |
| T=0.3, sample=38, score=0.7833, global_score=1.3323, seq_recovery=0.6923 | ASRLYTLVFVLQPQRVLLGMKKRGFGAGRWNGFGGKVQEGETIEDGARRELQEESGLTVDALHKVGQIVFEFVGEPELMDVHVFCTDSIQGTPVESDEMRPCWFQLDQIPFKDMWPDFSLWFPLLLQKKKFHGYFKFQGQDTILDYTLREVDTV |
| T=0.3, sample=39, score=0.7514, global_score=1.3456, seq_recovery=0.5385 | ASRLYTLVFCLQPQRVLLGMKKRGFGAGRWNGFGGKVQEGETIEDGARRELQEESGLTVDALHKVGQIVFEFVGEPELMDVHVFCTDSIQGTPVESDEMRPCWFQLDQIPFKNMFPDFSLWFPLLLQKKKFHGYFKFQGQDTILDYTLREVDTV |
| T=0.3, sample=40, score=0.7309, global_score=1.3337, seq_recovery=0.6154 | ASRLYTLVFCLQPQRVLLGMKKRGFGAGRWNGFGGKVQEGETIEDGARRELQEESGLTVDALHKVGQIVFEFVGEPELMDVHVFCTDSIQGTPVESDEMRPCWFQLDQIPFKQMHPDYSLWFPLLLQKKKFHGYFKFQGQDTILDYTLREVDTV |
| T=0.3, sample=41, score=0.7784, global_score=1.3570, seq_recovery=0.5385 | ASRLYTLVFCLQPQRVLLGMKKRGFGAGRWNGFGGKVQEGETIEDGARRELQEESGLTVDALHKVGQIVFEFVGEPELMDVHVFCTDSIQGTPVESDEMRPCWFQLDQIPFKEMFPDASLWFPLLLQKKKFHGYFKFQGQDTILDYTLREVDTV |
| T=0.3, sample=42, score=0.9068, global_score=1.3721, seq_recovery=0.4615 | ASRLYTLVFCLQPQRVLLGMKKRGFGAGRWNGFGGKVQEGETIEDGARRELQEESGLTVDALHKVGQIVFEFVGEPELMDVHVFCTDSIQGTPVESDEMRPCWFQLDQIPFKNMWPDFSLWYPLLLQKKKFHGYFKFQGQDTILDYTLREVDTV |
| T=0.3, sample=43, score=0.7654, global_score=1.3740, seq_recovery=0.6154 | ASRLYTLVFVLQPQRVLLGMKKRGFGAGRWNGFGGKVQEGETIEDGARRELQEESGLTVDALHKVGQIVFEFVGEPELMDVHVFCTDSIQGTPVESDEMRPCWFQLDQIPFKEMFPDASLWFPLLLQKKKFHGYFKFQGQDTILDYTLREVDTV |
| T=0.3, sample=44, score=0.7188, global_score=1.3642, seq_recovery=0.6154 | ASRLYTLVFCLQPQRVLLGMKKRGFGAGRWNGFGGKVQEGETIEDGARRELQEESGLTVDALHKVGQIVFEFVGEPELMDVHVFCTDSIQGTPVESDEMRPCWFQLDQIPFKDMYPDYSLWFPLLLQKKKFHGYFKFQGQDTILDYTLREVDTV |
| T=0.3, sample=46, score=0.7569, global_score=1.3481, seq_recovery=0.5385 | ASRLFTLVYCLQPQRVLLGMKKRGFGAGRWNGFGGKVQEGETIEDGARRELQEESGLTVDALHKVGQIVFEFVGEPELMDVHVFCTDSIQGTPVESDEMRPCWFQLDQIPFKDMFPDASLWFPLLLQKKKFHGYFKFQGQDTILDYTLREVDTV |
| T=0.3, sample=48, score=0.7767, global_score=1.3365, seq_recovery=0.6154 | ASRLYTLVYCLQPQRVLLGMKKRGFGAGRWNGFGGKVQEGETIEDGARRELQEESGLTVDALHKVGQIVFEFVGEPELMDVHVFCTDSIQGTPVESDEMRPCWFQLDQIPFKDMWPDASLWFPLLLQKKKFHGYFKFQGQDTILDYTLREVDTV |
| T=0.3, sample=49, score=0.7156, global_score=1.3289, seq_recovery=0.5385 | ASRLYTLVYCLQPQRVLLGMKKRGFGAGRWNGFGGKVQEGETIEDGARRELQEESGLTVDALHKVGQIVFEFVGEPELMDVHVFCTDSIQGTPVESDEMRPCWFQLDQIPFKQMFPDASLWFPLLLQKKKFHGYFKFQGQDTILDYTLREVDTV |
| T=0.3, sample=50, score=0.7333, global_score=1.3487, seq_recovery=0.6154 | ASRLYTLVFCLQPQRVLLGMKKRGFGAGRWNGFGGKVQEGETIEDGARRELQEESGLTVDALHKVGQIVFEFVGEPELMDVHVFCTDSIQGTPVESDEMRPCWFQLDQIPFKDMFPDYSLWFPLLLQKKKFHGYFKFQGQDTILDYTLREVDTV |
| T=0.3, sample=51, score=0.6906, global_score=1.3577, seq_recovery=0.6154 | ASRLYTLVFCLQPQRVLLGMKKRGFGAGRWNGFGGKVQEGETIEDGARRELQEESGLTVDALHKVGQIVFEFVGEPELMDVHVFCTDSIQGTPVESDEMRPCWFQLDQIPFKDMWPDASLWFPLLLQKKKFHGYFKFQGQDTILDYTLREVDTV |
| T=0.3, sample=52, score=0.8029, global_score=1.3596, seq_recovery=0.6923 | ASRLYTLVFCLQPQRVLLGMKKRGFGAGRWNGFGGKVQEGETIEDGARRELQEESGLTVDALHKVGQIVFEFVGEPELMDVHVFCTDSIQGTPVESDEMRPCWFQLDQIPFKDMYPDYSYWFPLLLQKKKFHGYFKFQGQDTILDYTLREVDTV |
| T=0.3, sample=53, score=0.7407, global_score=1.3483, seq_recovery=0.6923 | ASRLYTLVFVLQPQRVLLGMKKRGFGAGRWNGFGGKVQEGETIEDGARRELQEESGLTVDALHKVGQIVFEFVGEPELMDVHVFCTDSIQGTPVESDEMRPCWFQLDQIPFKDMWPDYSLWFPLLLQKKKFHGYFKFQGQDTILDYTLREVDTV |
| T=0.3, sample=54, score=0.8690, global_score=1.3855, seq_recovery=0.6154 | ASRLYTLVFALQPQRVLLGMKKRGFGAGRWNGFGGKVQEGETIEDGARRELQEESGLTVDALHKVGQIVFEFVGEPELMDVHVFCTDSIQGTPVESDEMRPCWFQLDQIPFKNMHPDYSLWFPLLLQKKKFHGYFKFQGQDTILDYTLREVDTV |
| T=0.3, sample=55, score=0.7989, global_score=1.3693, seq_recovery=0.6154 | ASRLYTLVFVLQPQRVLLGMKKRGFGAGRWNGFGGKVQEGETIEDGARRELQEESGLTVDALHKVGQIVFEFVGEPELMDVHVFCTDSIQGTPVESDEMRPCWFQLDQIPFKQMFPDASLWFPLLLQKKKFHGYFKFQGQDTILDYTLREVDTV |
| T=0.3, sample=56, score=0.7200, global_score=1.3697, seq_recovery=0.5385 | ASRLYTLVFCLQPQRVLLGMKKRGFGAGRWNGFGGKVQEGETIEDGARRELQEESGLTVDALHKVGQIVFEFVGEPELMDVHVFCTDSIQGTPVESDEMRPCWFQLDQIPFKQMFPDYSLWFPLLLQKKKFHGYFKFQGQDTILDYTLREVDTV |
| T=0.3, sample=57, score=0.7345, global_score=1.3420, seq_recovery=0.6154 | ASRLYTLVFCLQPQRVLLGMKKRGFGAGRWNGFGGKVQEGETIEDGARRELQEESGLTVDALHKVGQIVFEFVGEPELMDVHVFCTDSIQGTPVESDEMRPCWFQLDQIPFKDMWPDFSLWFPLLLQKKKFHGYFKFQGQDTILDYTLREVDTV |
| T=0.3, sample=60, score=0.7559, global_score=1.3441, seq_recovery=0.5385 | ASRLYTLVFCLQPQRVLLGMKKRGFGAGRWNGFGGKVQEGETIEDGARRELQEESGLTVDALHKVGQIVFEFVGEPELMDVHVFCTDSIQGTPVESDEMRPCWFQLDQIPFKNMFPDASLWFPLLLQKKKFHGYFKFQGQDTILDYTLREVDTV |
| T=0.3, sample=61, score=0.7295, global_score=1.3801, seq_recovery=0.5385 | ASRLYTLVFCLQPQRVLLGMKKRGFGAGRWNGFGGKVQEGETIEDGARRELQEESGLTVDALHKVGQIVFEFVGEPELMDVHVFCTDSIQGTPVESDEMRPCWFQLDQIPFKQMWPDFSLWFPLLLQKKKFHGYFKFQGQDTILDYTLREVDTV |
| T=0.3, sample=63, score=0.7445, global_score=1.3339, seq_recovery=0.5385 | ASRLYTLVFCLQPQRVLLGMKKRGFGAGRWNGFGGKVQEGETIEDGARRELQEESGLTVDALHKVGQIVFEFVGEPELMDVHVFCTDSIQGTPVESDEMRPCWFQLDQIPFKEMFPDFSLWFPLLLQKKKFHGYFKFQGQDTILDYTLREVDTV |
| T=0.3, sample=64, score=0.7614, global_score=1.3390, seq_recovery=0.5385 | ASRLFTLVFCLQPQRVLLGMKKRGFGAGRWNGFGGKVQEGETIEDGARRELQEESGLTVDALHKVGQIVFEFVGEPELMDVHVFCTDSIQGTPVESDEMRPCWFQLDQIPFKDMYPDYSLWFPLLLQKKKFHGYFKFQGQDTILDYTLREVDTV |
| T=0.3, sample=65, score=0.7989, global_score=1.3585, seq_recovery=0.6154 | ASRLYTLVFVLQPQRVLLGMKKRGFGAGRWNGFGGKVQEGETIEDGARRELQEESGLTVDALHKVGQIVFEFVGEPELMDVHVFCTDSIQGTPVESDEMRPCWFQLDQIPFKNMFPDYSLWFPLLLQKKKFHGYFKFQGQDTILDYTLREVDTV |
| T=0.3, sample=66, score=0.7515, global_score=1.3537, seq_recovery=0.5385 | ASRLYTLVFCLQPQRVLLGMKKRGFGAGRWNGFGGKVQEGETIEDGARRELQEESGLTVDALHKVGQIVFEFVGEPELMDVHVFCTDSIQGTPVESDEMRPCWFQLDQIPFKEMWPDYSLWFPLLLQKKKFHGYFKFQGQDTILDYTLREVDTV |
| T=0.3, sample=69, score=0.7697, global_score=1.3681, seq_recovery=0.6923 | ASRLYTLVFVLQPQRVLLGMKKRGFGAGRWNGFGGKVQEGETIEDGARRELQEESGLTVDALHKVGQIVFEFVGEPELMDVHVFCTDSIQGTPVESDEMRPCWFQLDQIPFKEMHPDFSLWFPLLLQKKKFHGYFKFQGQDTILDYTLREVDTV |
| T=0.3, sample=71, score=0.7712, global_score=1.3562, seq_recovery=0.6923 | ASRLYTLVFCLQPQRVLLGMKKRGFGAGRWNGFGGKVQEGETIEDGARRELQEESGLTVDALHKVGQIVFEFVGEPELMDVHVFCTDSIQGTPVESDEMRPCWFQLDQIPFKDMHPDYSLWFPLLLQKKKFHGYFKFQGQDTILDYTLREVDTV |
| T=0.3, sample=75, score=0.7225, global_score=1.3530, seq_recovery=0.4615 | ASRLFTLVYCLQPQRVLLGMKKRGFGAGRWNGFGGKVQEGETIEDGARRELQEESGLTVDALHKVGQIVFEFVGEPELMDVHVFCTDSIQGTPVESDEMRPCWFQLDQIPFKQMWPDASLWFPLLLQKKKFHGYFKFQGQDTILDYTLREVDTV |
| T=0.3, sample=77, score=0.7738, global_score=1.3331, seq_recovery=0.5385 | ASRLFTLVFCLQPQRVLLGMKKRGFGAGRWNGFGGKVQEGETIEDGARRELQEESGLTVDALHKVGQIVFEFVGEPELMDVHVFCTDSIQGTPVESDEMRPCWFQLDQIPFKEMHPDFSLWFPLLLQKKKFHGYFKFQGQDTILDYTLREVDTV |
| T=0.3, sample=82, score=0.7591, global_score=1.3623, seq_recovery=0.5385 | ASRLYTLVFCLQPQRVLLGMKKRGFGAGRWNGFGGKVQEGETIEDGARRELQEESGLTVDALHKVGQIVFEFVGEPELMDVHVFCTDSIQGTPVESDEMRPCWFQLDQIPFKNMFPDYSLWFPLLLQKKKFHGYFKFQGQDTILDYTLREVDTV |
| T=0.3, sample=85, score=0.7804, global_score=1.3340, seq_recovery=0.6154 | ASRLYTLVYCLQPQRVLLGMKKRGFGAGRWNGFGGKVQEGETIEDGARRELQEESGLTVDALHKVGQIVFEFVGEPELMDVHVFCTDSIQGTPVESDEMRPCWFQLDQIPFKNMHPDASLWFPLLLQKKKFHGYFKFQGQDTILDYTLREVDTV |
| T=0.3, sample=89, score=0.7378, global_score=1.3694, seq_recovery=0.5385 | ASRLYTLVFCLQPQRVLLGMKKRGFGAGRWNGFGGKVQEGETIEDGARRELQEESGLTVDALHKVGQIVFEFVGEPELMDVHVFCTDSIQGTPVESDEMRPCWFQLDQIPFKQMFPDFSLWFPLLLQKKKFHGYFKFQGQDTILDYTLREVDTV |
| T=0.3, sample=90, score=0.7380, global_score=1.3437, seq_recovery=0.6154 | ASRLYTLVFCLQPQRVLLGMKKRGFGAGRWNGFGGKVQEGETIEDGARRELQEESGLTVDALHKVGQIVFEFVGEPELMDVHVFCTDSIQGTPVESDEMRPCWFQLDQIPFKDMYPDFSLWFPLLLQKKKFHGYFKFQGQDTILDYTLREVDTV |
| T=0.3, sample=95, score=0.8117, global_score=1.3659, seq_recovery=0.4615 | ASRLFTLVFCLQPQRVLLGMKKRGFGAGRWNGFGGKVQEGETIEDGARRELQEESGLTVDALHKVGQIVFEFVGEPELMDVHVFCTDSIQGTPVESDEMRPCWFQLDQIPFKNMYPDFSLWFPLLLQKKKFHGYFKFQGQDTILDYTLREVDTV |
| T=0.3, sample=97, score=0.8266, global_score=1.3401, seq_recovery=0.5385 | ASRLYTLVFCLQPQRVLLGMKKRGFGAGRWNGFGGKVQEGETIEDGARRELQEESGLTVDALHKVGQIVFEFVGEPELMDVHVFCTDSIQGTPVESDEMRPCWFQLDQIPFKEMNPDFSLWFPLLLQKKKFHGYFKFQGQDTILDYTLREVDTV |
